# A selective and rapid cell-permeable inhibitor of human caspase-3

**DOI:** 10.1101/706614

**Authors:** Angelo Solania, Gonzalo E. González-Páez, Dennis W. Wolan

**Affiliations:** Departments of Molecular Medicine and Integrative Structural and Computational Biology, The Scripps Research Institute, 10550 North Torrey Pines Road, La Jolla, CA 92037

**Keywords:** Caspase-3, peptide-based inhibitor, unnatural amino acids, cell-permeable

## Abstract

The individual roles and overlapping functionalities the twelve human caspases have during apoptosis and other cellular processes remain poorly resolved primarily due to a lack of chemical tools. Here we present a new selective caspase-3 inhibitor, termed Ac-ATS010-KE, with rapid and irreversible binding kinetics. Relative to previously designed caspase-3-selective molecules that have tremendously abated inhibitory rates and thus limited use in biological settings, the improved kinetics of Ac-ATS010-KE permit its use in a cell-based capacity. We demonstrate that Ac-ATS010-KE prevents apoptosis with comparable efficacy to the general caspase inhibitor Ac-DEVD-KE and surprisingly does so without side-chain methylation. This observation is in contrast to the well-established peptide modification strategy typically employed for improving cellular permeability. Ac-ATS010-KE protects against extrinsic apoptosis, which demonstrates the utility of a thiophene carboxylate leaving group in biological settings, challenges the requisite neutralization of free carboxylic acids to improve cell permeability, and provides a tool-like compound to interrogate the role of caspase-3 in a variety of cellular processes.

---

The essentiality of executioner caspase-3 (casp-3) to the final stages of apoptosis and the cellular substrates targeted by this cysteine protease remains a fundamental question since its discovery over 30 years ago. While the consensus suggests casp-3 is critical, some studies have demonstrated casp-3 to be nonessential to apoptosis. For example, casp-3-null MCF-7 cells respond to apoptotic stimuli^1^ and may employ caspase-independent pathways^2, 3^ and/or compensatory overexpression of other caspases to advance apoptosis. Peptide-based substrates and inhibitors are routinely used in cell-based assays to detect or prevent caspase activity, respectively; however, these chemical tools are too promiscuous to confidently attribute results to individual caspases within normal cellular contexts.^4^ For example, casp-3 and the highly homologous executioner caspase-7 (casp-7) have identical *in vitro* peptide substrate profiles with the optimal sequence of the tetrapeptide DEVD.^5^

N-terminal labeling in combination with mass-spectrometry degradomic analyses of cellular lysates confirmed the canonical sequence preferences originally established using *in vitro* fluorogenic substrate libraries. Additionally, these studies have led to the identification of thousands of apoptotic caspase substrates.^6–8^ Unfortunately, these studies could not resolve the protease(s) responsible for the cleavage events. Recent cell-free lysate studies and the divergent phenotypes observed between casp-3 and casp-7 knockout mouse models indicate the strong likelihood that these proteases have distinct cellular substrates.^9, 10^ This disconnect between identical *in vitro* specificities of casp-3 and casp-7 with the cell-based and *in vivo* studies may be partially explained by potential exosite interactions between enzymes and their full length biological substrates.^11^ While mass spectrometry based studies have identified differences in the prime side preferences of casp-3 and casp-7 (e.g., C-terminal residues with respect to the site of hydrolysis),^12^ probes that utilize this prime-side information have yet to be designed.

The development of unnatural amino acid-containing caspase-specific substrates that allow for the selective detection of an individual caspase’s activity has recently been reported.^4, 5, 13–15^ Notwithstanding, these tools do not enable us to regulate and thus deconvolute the biologically relevant substrates and mechanisms of specific caspases. This endeavor will only be possible with potent, selective, and universally applicable cell-permeable inhibitors. A casp-3-selective molecule with dramatically improved spatial and temporal resolution would help define the protease’s role across a range of cell types. In addition, compounds like this can be applied in dose- and time-dependent experiments unlike genetic manipulation, which are typically restricted to endpoint assays only. The current standard for chemically regulating caspase activity are commercially available fluoromethyl ketone (FMK) “prodrug” inhibitors, such as Z-VAD^M^-FMK and Z-D^M^E^M^VD^M^-FMK (^M^ denotes O-methylation of side chain). These peptides rely on general cellular esterases to unmask the O-methyl esters into carboxylates for optimal caspase binding to protect against cell death by potently inhibiting several caspases.^15, 16^

We previously identified a 5-residue peptide sequence containing unnatural amino acids selective for casp-3.^17^, ^18^ Our peptide resolved the activity of casp-3 in apoptotic cells as a fluorescence quenched cell-permeable substrate and a probe to detect active caspase-3. Unfortunately, appending a unique 5-methyl-2-thiophene carboxylic acid (KE) leaving group onto the C-terminus to generate a selective casp-3 irreversible competitive inhibitor, termed Ac-DW3-KE (**8**), yielded a molecule unable to protect against apoptotic cell death.^18^ The inability of **8** to protect against apoptosis may suggest that casp-3 is dispensable to apoptotic cell death, as observed for MCF-7 cells. However, other factors may have accounted for the lack of activity, including: 1) the potential of cellular toxicity of the displaced KE warhead upon reacting with casp-3; 2) a decreased rate and/or limited amount in peptide cellular internalization; 3) insufficient rates of casp-3 inhibition with respect to caspase activation during apoptosis; and 4) a reduced compatibility with general esterase activity required to de-esterify the O-methylated peptide “prodrug” into its active form. Thus, we sought to explore if these key variables separately, or in combination, limit the application of **8** in cellular studies.^18^

Here, we present an optimized casp-3-selective inhibitor, Ac-ATS010-KE (**10**), that rapidly and selectively inhibits casp-3 in a cellular setting and protects cells against extrinsically induced apoptosis. Furthermore, the apoptotic protection afforded by **10** is improved in the absence of side chain methylation. This result suggests that cellular permeability may not be the limiting factor for these peptide-based inhibitors, which challenges the use of side chain O-methylation that has been ubiquitously applied to caspase inhibitors since the early observations that O-methylation of the P1 Asp improved the stability of Z-VAD-FMK.^19^

## Results and Discussion

### Peptides containing 5-methyl-2-thiophene carboxylate leaving group are useful for biological studies

Three activated methyl ketone inhibitors: Z-D^M^E^M^VD^M^-FMK (**0^M^**), Ac-D^M^E^M^VD^M^-KE (**1^M^**), and Ac-DW3^M^-KE (**8^M^**) were evaluated to determine their ability to prevent cell death due to extrinsically induced apoptosis (Figure 1A). Acute T cell lymphoblasts (Jurkat cells) were pretreated with 20 μM of the indicated inhibitor for 3 h before induction of extrinsic apoptosis with 10 ng/mL MegaFasL. Cellular viability was then quantified using CellTiter-Glo^®^. Treatment with the KE-containing **1^M^** peptide protected Jurkat cells from apoptosis better than the commercially available **0^M^**. Thus, the 5-methyl-2-thiophene carboxylate is a viable warhead for cellular studies and does not promote cell death upon release (Figure 1B).

**Figure 1.**
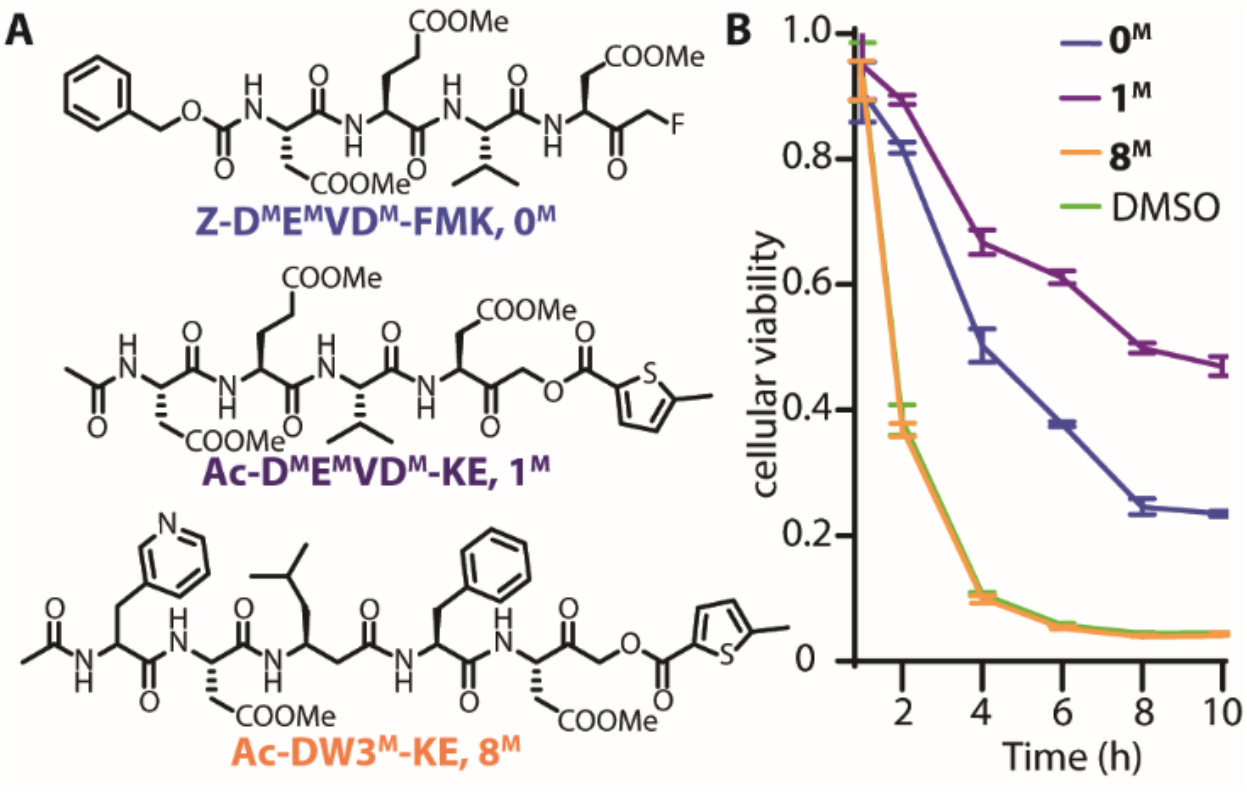
Ac-DW3^**M**^KE (**8^M^**) is not suitable for cellular studies. (A) Chemical structures of promiscuous, commercially available Z-D^**M**^E^**M**^VD^**M**^-FMK (**0^M^**), a KE derivative (**1^M^**), and casp-3-selective inhibitor (**8^M^**). (B) Extrinsic apoptosis protection assay. Jurkat cells were treated for 3 h with 20 μM inhibitor before apoptosis was induced with 10 ng/mL MegaFasL. At the indicated time cellular viability was measured using CellTiter-Glo®. Pan-caspase inhibitors (**0^M^** and **1^M^**) were able to preserve cellular viability and casp-3-selective inhibitor (**8^M^**) was not. The 5-methyl-2-thiophene carboxylate (KE) is an effective leaving group for cellular studies.

### Ac-DW3-KE is a slow *in vitro* caspase-3 inhibitor

*In vitro* kinetic assays of **8** and related peptide inhibitors based on our previously published DW3 sequence revealed that the combination of a P3 ß-homoleucine (ßhLeu) with a P2 Phe (e.g., **4** and **8**) had prohibitively poor binding kinetics with respect to the canonical DEVD sequence in **1** (Table 1). Critically, the inactivation of casp-3 by **8** was 500x less efficient than the inactivation of casp-3 by **1**. These results suggest that inhibitors containing the recognition sequence of **8** are not viable for cellular studies, as the rate of inhibition would be likely superseded by the rate of casp-3 activation and promotion of downstream apoptotic events. Future inhibitors will need to be more selective and as potent as DEVD-based inhibitors which possess low nanomolar/high picamolar affinity toward casp-3. We posit, an inhibitor is sufficiently potent if the molecule can preserve cellular viability after apoptotic induction, as observed for both compounds **0^M^** and **1^M^** (Figure 1B).

**Table 1.** Kinetic efficiencies and selectivities of DW3 inhibitor sub-library.

| cmpd | Ac-Sequence-KE |  |  |  |  | K<sub>i</sub> (μM) |  |  |  | k<sub>inact</sub> (× 10<sup>-3</sup> s<sup>-1</sup>) |  |  |  | k<sub>inact</sub>/K<sub>i</sub> (× 10<sup>4</sup> s<sup>-1</sup> M<sup>-1</sup>) |  |  |  | casp-3 selectivity |  |  |
| --- | --- | --- | --- | --- | --- | --- | --- | --- | --- | --- | --- | --- | --- | --- | --- | --- | --- | --- | --- | --- |
|  | P5 | P4 | P3 | P2 | P1 | caspase |  |  |  | caspase |  |  |  | caspase |  |  |  | /casp-6 | /casp-7 | /casp-8 |
|  |  |  |  |  |  | -3 | -6 | -7 | -8 | -3 | -6 | -7 | -8 | -3 | -6 | -7 | -8 |  |  |  |
| 1 |  | D | E | V | D | 0.0012 | 0.029 | 0.012 | 0.013 | 6.9 | 7.2 | 7.3 | 6.5 | 550 | 25 | 60 | 51 | 22 | 9.2 | 11 |
| 2 |  | D | E | F | D | 0.0028 | 0.13 | 0.016 | 0.0041 | 4.8 | 4.0 | 5.8 | 2.6 | 170 | 3.2 | 36 | 64 | 54 | 4.7 | 2.7 |
| 3 |  | D | βhLeu | V | D | 0.048 | 0.40 | 0.78 | 0.31 | 5.1 | 2.9 | 5.8 | 2.8 | 11 | 0.74 | 0.75 | 0.92 | 14 | 14 | 12 |
| 4 |  | D | βhLeu | F | D | 1.3 | 2.8 | 25 | 18 | 5.0 | 3.7 | 6.3 | 9.0 | 0.38 | 0.13 | 0.025 | 0.051 | 2.8 | 15 | 7.4 |
| 5 | 3PAI | D | E | V | D | 0.0081 | 0.05 | 0.049 | 0.022 | 6.2 | 4.3 | 4.9 | 3.4 | 77 | 8.1 | 9.9 | 15 | 9.5 | 7.8 | 5.1 |
| 6 | 3PAI | D | E | F | D | 0.0025 | 0.05 | 0.019 | 0.025 | 9.4 | 6.7 | 7.7 | 5.7 | 380 | 13 | 41 | 23 | 29 | 9.2 | 17 |
| 7 | 3PAI | D | βhLeu | V | D | 0.059 | 1.1 | 0.78 | 0.27 | 4.8 | 3.4 | 4.3 | 2.7 | 8.0 | 0.30 | 0.55 | 1.0 | 26 | 15 | 8.0 |
| 8 | 3PAI | D | βhLeu | F | D | 0.49 | 1.0 | 9.6 | 3.0 | 5.3 | 1.5 | 5.4 | 3.7 | 1.1 | 0.16 | 0.056 | 0.12 | 6.9 | 19 | 8.6 |

### Discovery and validation of a potent casp-3-selective inhibitor

Due to the slow binding kinetics of **8**, we replaced the P3 ßhLeu with a pentofluorophenylalanine (PF5), as this residue was previously shown to impart selectivity against casp-8.^13^ The resulting molecule, **10**, had a k_inact_/K_i_ for casp-3 5480x, 9x, 68x, 121x, 1700x higher over casp-6, casp-7, casp-8, casp-9 and casp-10, respectively (Table 2).

**Table 2.** Kinetic efficiencies of potent caspase-3-selective and related inhibitors.

| Sequence |  |  |  |  |  |  |  | K<sub>i</sub> (μM) |  |  |  |  |  | K<sub>inact</sub> (× 10<sup>-3</sup> s<sup>-1</sup>) |  |  |  |  |  | K<sub>inact</sub>/K<sub>i</sub> (× 10<sup>4</sup> s<sup>-1</sup> M<sup>-1</sup>) |  |  |  |  |  |
| --- | --- | --- | --- | --- | --- | --- | --- | --- | --- | --- | --- | --- | --- | --- | --- | --- | --- | --- | --- | --- | --- | --- | --- | --- | --- |
| cmpd | N-Cap | P5 | P4 | P3 | P2 | P1 | War head | caspase |  |  |  |  |  | caspase |  |  |  |  |  | caspase |  |  |  |  |  |
|  |  |  |  |  |  |  |  | -3 | -6 | -7 | -8 | -9 | -10 | -3 | -6 | -7 | -8 | -9 | -10 | -3 | -6 | -7 | -8 | -9 | -10 |
| 9<sup>a</sup> | Ac |  | D | E | V | D | CHO | 0.00023 | 0.031 | 0.0016 | 0.00092 | 0.060 | 0.012 |  |  |  |  |  |  |  |  |  |  |  |  |
| 1<sup>M</sup> | Ac |  | D<sup>M</sup> | E<sup>M</sup> | V<sup>M</sup> | D<sup>M</sup> | KE | 0.29 | 3.2 | 1.5 | 0.22 | 2.3 | 2.0 | 8.3 | 1.4 | 7.6 | 1.0 | 4.1 | 1.0 | 2.9 | 0.043 | 0.52 | 0.44 | 0.17 | 0.052 |
| 1 | Ac |  | D | E | V | D | KE | 0.0012 | 0.029 | 0.012 | 0.013 | 0.11 | 0.048 | 6.9 | 7.2 | 7.3 | 6.5 | 4.2 | 3.7 | 560 | 25 | 60 | 51 | 3.7 | 7.7 |
| B0 | Bio |  | D | E | V | D | KE | 0.0012 | 0.018 | 0.014 | 0.020 | 0.17 | 0.090 | 7.5 | 7.1 | 8.5 | 6.5 | 6.4 | 4.8 | 610 | 39 | 61 | 32 | 3.8 | 5.3 |
| 6 | Ac | 3PAI | D | E | F | D | KE | 0.0025 | 0.052 | 0.019 | 0.025 | 0.18 | 0.16 | 9.4 | 6.7 | 7.7 | 5.7 | 5.4 | 4.2 | 380 | 13 | 41 | 23 | 2.9 | 2.7 |
| B6 | Bio | 3PAI | D | E | F | D | KE | 0.0022 | 0.051 | 0.018 | 0.025 | 0.22 | 0.20 | 10. | 6.9 | 9.1 | 6.1 | 7.2 | 4.3 | 480 | 14 | 51 | 25 | 3.3 | 2.1 |
| 10 | Ac | 3PAI | D | P5F | F | D | KE | 0.0059 | 14 | 0.045 | 0.20 | 0.28 | 1.8 | 10. | 4.3 | 8.3 | 5.1 | 3.9 | 1.8 | 170 | 0.031 | 19 | 2.5 | 1.4 | 0.10 |
| B10 | Bio | 3PAI | D | P5F | F | D | KE | 0.0040 | 10. | 0.093 | 0.92 | 0.53 | 0.23 | 2.2 | 9.6 | 4.2 | 5.5 | 4.6 | 0.30 | 55 | 0.094 | 4.5 | 0.60 | 0.86 | 0.15 |
| 11 | Ac | 3PAI | D | βhLeu | hLeu | D | KE | 0.98 | 23 | 49 | .82 | 3.4 | 1.9 | 3.0 | 0.9 | 1.6 | 2.0 | 0.86 | 2.5 | 3.0 | 0.039 | 0.033 | 2.4 | 0.25 | 1.29 |

<sup>a</sup> values referenced from (Thornberry et al., 1998); <sup>M</sup> denotes an O-methylated sidechain

Importantly, conversion of the optimized inhibitors into the corresponding probe by incorporation of an N-terminal d-biotin (**B10**) had marginal effect on the potency and selectivity of the compounds (Table 2). This conservation in potency was anticipated, as structures of casp-3 in complex with various peptide inhibitors show that residues beyond the P5 position lie outside the active site of casp-3 and consequently have limited direct interaction with the protease. We envision these probes will be enable us to enrich and detect active caspases in non-apoptotic processes.

### Structural analysis of casp-3 and casp-7 in complex with inhibitors

To establish how this molecule generates selectivity and potentially guide future iterations of casp-3-selective inhibitors, the co-crystal structures of human casp-3 and casp-7 in complex with **8** and **10** were determined. These structures were compared to the co-crystal structures of Ac-3PAl-D-ßhLeu-hLeu-D-KE (**11**), our first generation casp-3-selective inhibitor, bound to casp-3 and casp-7 (3-Pal: 3-pyridylalanine; hLeu: homoleucine).^17^ As we have previously shown, the additional P5 3PAl residue on **8**, **10**, and **11** forms two additional interactions with S209 of casp-3 relative to casp-7 (Figure 2A).^17^ No equivalent interactions are observed in the casp-7:**10** co-crystal structure, as casp-7 has a proline in place of S209 (Figure 2B). These additional hydrogen bonds are highlighted in Figure 2C (red dashed lines) and all structure interaction schematics for casp-7:**8**, casp-7:**10**, and casp-3:**8** are provided in Figure S2.

**Figure 2.**
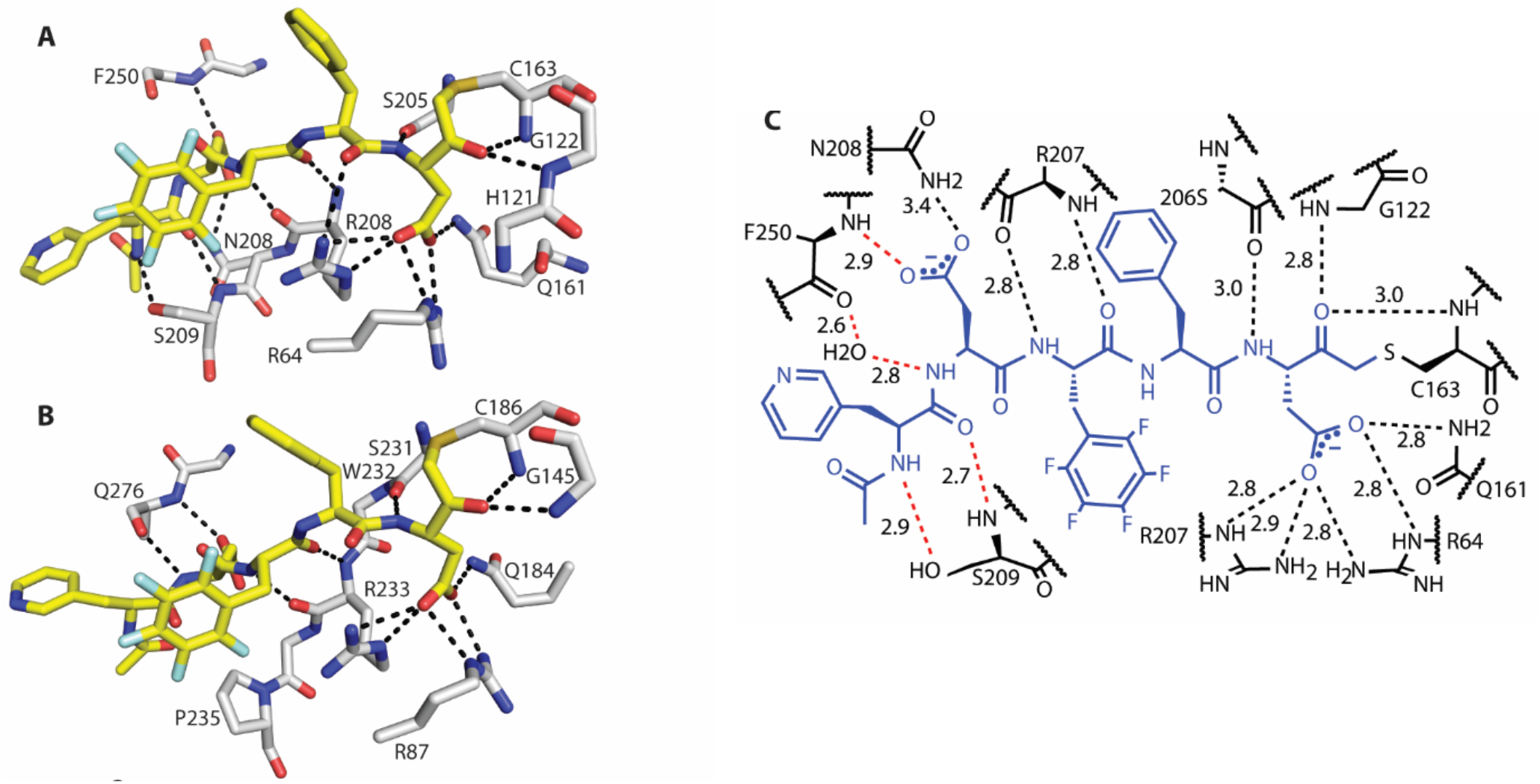
Structural characterization of co-crystal structures with casp-3-selective inhibitor **10** using x-ray crystallography. (A) Co-crystal structure (PDB ID:6CL0) of casp-3 (gray):**10** (yellow) highlighting critical hydrogen bonds. (B) Co-crystal structure of casp-7 (gray):**10** (yellow) (PDB ID:6CL2). (C) Schematic of casp-3:**10** interactions depicting unique hydrogen bonds (red) relative to the casp-7:**10**.

Superposition with a Z-DEVD-bound structure (PDB ID: 2DKO) shows that **8** and **11** adopt significant energetically unfavorable backbone confirmations to preserve optimal side chain interactions within the casp-3 active site (Figures 3A and 3B). Specifically, casp-3 accomodates the additional P3 backbone methylene of **8** and **11** while maintaining the positions of the P1 and P4 aspartic acid side chains. This is accomplished by forcing the extended peptide backbone of **8** and **11** to kink with unfavorable phi/psi angles to fit within the same space (Figures 3A and B). Furthermore, a key hydrogen bond between the P3 amide nitrogen of DEVD and the casp-3 R207 main-chain carbonyl is eliminated in the casp-3:**8** co-crystal structure (Figure 3A). The **8**-co-crystal structure provides a strong rationalization for the poor inactivation kinetics (Table 1, Figure 3A).

**Figure 3.**
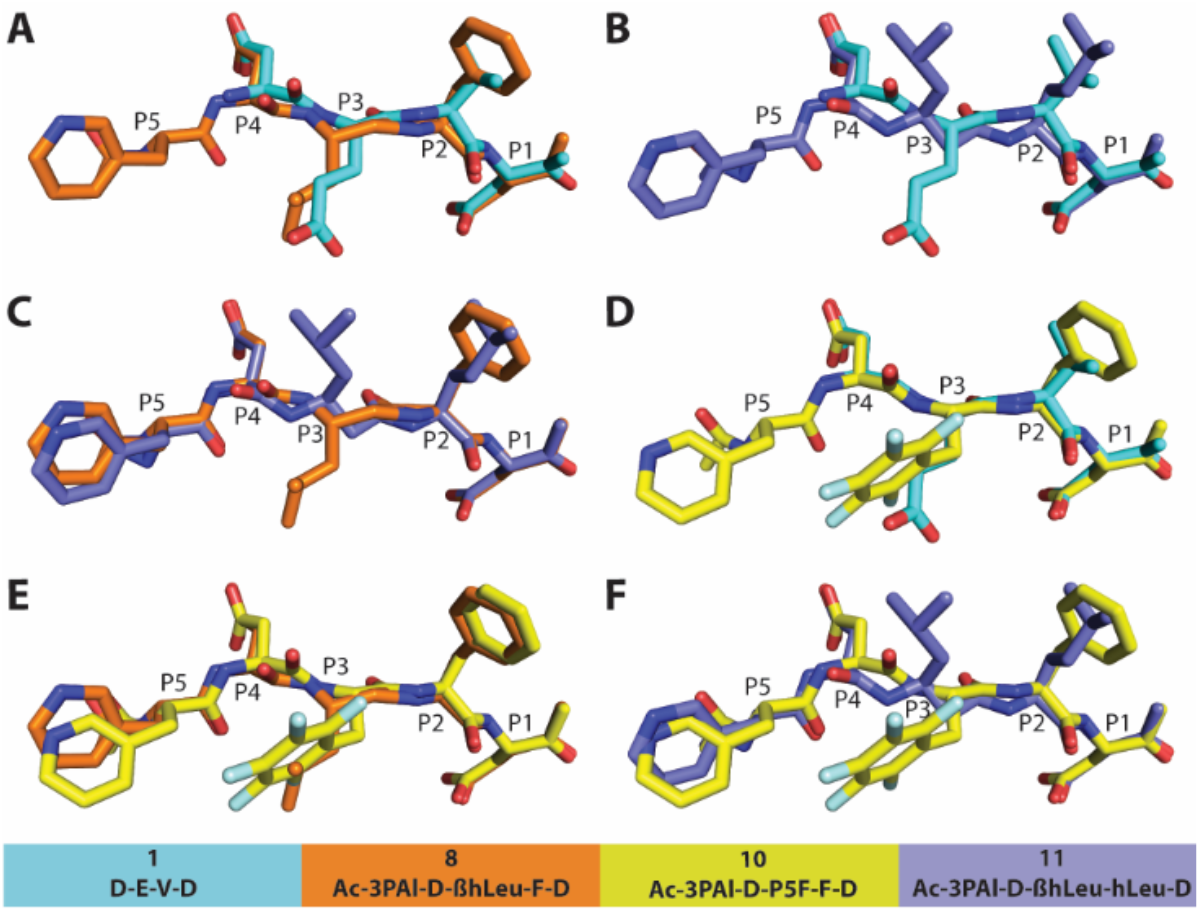
Structural comparisons of **1** (PDB ID: 2KDO), **8** (PDB ID: 6CKZ), **10** (PDB ID: 6CL0), and **11** (PDB ID:4JJE) bound to casp-3. (A) Overlay of **1** and **8** demonstrating the perturbed backbone conformation of **8**. (B) Overlay of **1** and **11** demonstrating the perturbed backbone confirmation of **11**. (C) Overlay of **8** and **11** demonstrating the high flexibility of the P3 ßhLeu. (D) Overlay demonstrating **1** and **10** bind with high conformational similarity. Overlay of **8** (E) and **11** (F) with **10** demonstrate older generation casp-3-selective inhibitors bind very differently compared to **10**.

Surprisingly, in comparison to **1** and **8**, the P3 ßhLeu side chain of **11** is flipped almost 180° with respect to the plane of the peptide backbone (Figure 3C). The observed flexibility of the P3 sidechain in the active site of casp-3 further demonstrates the poor fit of the P3 ßhLeu and consequently, the slow kinetics of inhibitors bearing this unnatural amino acid toward casp-3. We hypothesize that this backbone elongation causes more severe perturbations in casp-7, as crystals of casp-7 bound with **8** fail to show conclusive density for the ligand past the P2 position (Figure S1). This observation strongly supports the poor binding affinity of this ligand to casp-7.

Our optimized casp-3-selective inhibitor **10** eliminates the elongated peptide backbone of our first-generation casp-3 molecules by substituting the P3 ßhLeu for PF5. Removal of the additional ß-methylene eliminates the corresponding unfavorable torsional angles required for the ßhLeu-containing **8** and **11** inhibitors to bind casp-3 (Figures 3A and 3B, respectively). An overlay of **10** and DEVD bound to casp-3 reveals that the backbone conformations superimpose with limited deviations despite **10** being comprised of multiple unnatural side chains (Figure 3D). The structural comparisons of the ßhLeu-containing peptides **8** and **11** with **10** clearly show that the casp-3 active site is optimally configured for interaction with standard amino acid main chains (Figure 3D, 3E, and 3F). Thus, all future casp-3-selective peptide optimizations would likely be limited to standard main-chain configurations for rapid casp-3 binding, while the side chains can be further modified for improved affinity.

Non-O-methylated inhibitors protect cells from apoptosis. Historically, protease-targeted peptidyl inhibitors were prepared for cellular studies by converting the free side-chain carboxylic acids into O-methyl esters (e.g., per-O-methylation). This common practice was found to improve the stability and cellular permeability of the peptide inhibitors, thereby improving their use for *in cellulo* and *in vivo* applications.^19, 20^

Once inside the cell, these inactive “prodrug”-like peptides are “activated” by endogenous cytoplasmic general esterases that de-esterify and unmask the carboxylic acids. This practice is ubiquitous in the design and application of caspase inhibitors for cell-based studies due the preferences of caspases to recognize and prefer negatively charged residues in the P1, P3, or P4 position. Notably, recent studies have demonstrated the ability of caspases to recognize P1 glutamic acid and phosphoserine in addition to aspartic acid.^21^ Critically, some commercial suppliers fail to specify the presence or absence of O-methylation and market caspase peptide inhibitors as cell-permeable inhibitors of apoptosis.

To investigate the requirement for per-O-methylation, both non-O-methylated and per-O-methylated versions of **1**, **8**, **10**, and **11** were analyzed using apoptotic protection assays. Of the per-O-methylated inhibitors, only inhibitors bearing the DEVD recognition sequence protected against MegaFasL-induced extrinsic apoptosis (Figures 1A and 4A). Surprisingly, we observed that non-O-methylated inhibitors outperformed their respective per-O-methylated counterparts under identical experimental conditions (Figure 4B). Most significant is the inability of **10^M^** to prevent cell death up to concentrations of 100 μM (Figure 4A). Conversely, non-O-methylated **10** was nearly as effective as the non-O-methylated DEVD-based general caspase inhibitor **1** (Figure 4B). To verify that the preservation of cellular viability was the result of inhibiting apoptosis, cleavage of PARP was analyzed by Western blot (Figure 4E). These findings suggest that per-O-methylation of the side-chain carboxylic acids is likely unnecessary for membrane permeability and/or active transport into cells and should be considered in the design and optimization of peptide-based protease inhibitors for cellular studies. Importantly, the use of unnatural amino acids in the recognition sequences may ablate the ability of intracellular esterases to process O-methylated molecules and account for the lack of observed cellular protection.

**Figure 4.**
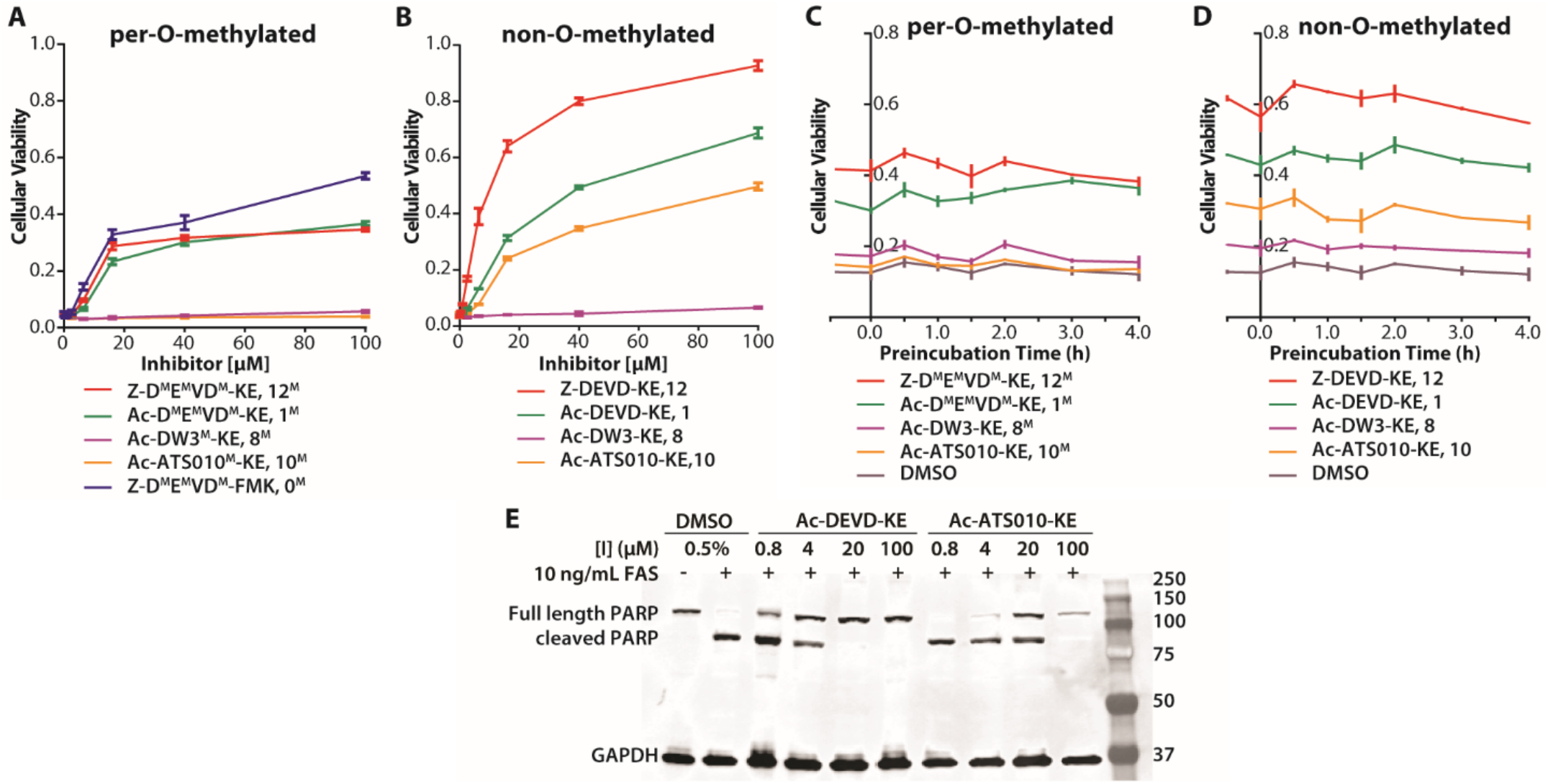
Evaluation of per-O-methylated and non-O-methylated caspase inhibitors in apoptosis protection assays. Extrinsic apoptosis was induced using 10 ng/mL MegaFasL. Cellular viability was measured using CellTiter-Glo^®^. (A & B) Jurkat cells were treated with the indicated concentration of inhibitor for 30 min before apoptosis was induced. Response curves of per-O-methylated inhibitors plateau similarly. Response curves of non-O-methylated inhibitors had a wider range and, in some cases, higher efficacy relative to the corresponding per-O-methylated “prodrug” version. (C & D) Jurkat cells were treated with 20 μM compound for the indicated time before apoptosis was induced and the amount of protection against apoptosis is not dependent on duration of pretreatment. (E) Western blot analysis of PARP cleavage and GAPDH in apoptotic Jurkat cells treated with **1**, **10**, or DMSO.

### O-methylation minimally affects inhibitor rate of internalization

To further investigate the utility of per-O-methylation, we performed a preincubation timecourse to assess differences in the rate of internalization of per-O-methylated versus non-O-methylated inhibitors. Jurkat cells were treated with 20 μM of the indicated inhibitor at various timepoints before and after apoptotic induction. Cells were harvested 3 h post MegaFasL addition and subjected to CellTiter-Glo^®^ analysis to quantify cellular viability. To our surprise, both non-O- and per-O-methylated inhibitors prevented cell death regardless of the duration of pre-incubation (Figure 4C and 4D). Furthermore, the addition of non-O-methylated or per-O-methylated inhibitors 30 min after apoptotic induction still protected against apoptosis. This phenomenon suggests that, at least for our purposes, both non-O-methylated and per-O-methylated inhibitors enter cells rapidly and cellular esterases act quickly to unmask the “prodrug” versions.

### Casp-3 inhibition alters caspase maturation during apoptosis

Apoptosis begins with the activation of casp-8 (extrinsic) or casp-9 (intrinsic) and both initiator caspases converge to activate casp-3, casp-6, and casp-7. We sought to determine if **10** protects cells against apoptosis similarly to the general caspase inhibitor **1** with suppression of caspase activation. Jurkat cells, pre-incubated with 20 μM **1** or **10** were collected hourly for 5 h post MegaFasL apoptosis induction and lysed by freeze-thaw. Lysates were subjected to Western blot analysis to quantitate activation of caspases-3, −6, −7, −8, and- 9 (Figure 5). As expected, caspase activation was completed within 2 h of apoptotic induction in Jurkat cells treated with the DMSO vehicle (Figure 5, lanes 1-4). We hypothesized that due to the promiscuity of **1** and potency against casp-8 (as measured by *in vitro* assays),**^18^** protection from extrinsic apoptosis was due to inactivation of upstream initiator casp-8, which precludes the activation of executioner casp-3 and casp-7. Indeed, the presence of the pan-inhibitor **1** resulted in the general suppression of caspase activation and is best exemplified by the persistence of the inactive proforms of casp-3 and casp-7 over 5 h (Figure 5, lanes 5-9). Unlike **1**, 20 μM **10** did not suppress caspase activation of all the apoptotic caspases. In fact, the caspase activation profile in the presence of **10** generally mimicked the DMSO-treated cells with the disappearance of proforms of casp-6, casp-7, casp-8, casp-9 (and appearance of active forms) occuring within the first 2 h after induction of apoptosis (Figure 5, lanes 10-14). However, one exceptional difference observed, is the persistance of the p20 large domain of casp-3 during cell death. In the absence of any inhibitor, the casp-3 large domain fully matures into the p17 form. This 3 kDa difference between p17 and p20 is due to the removal of the N-terminal peptide that indicates complete casp-3 maturation (Figure 5, lanes 10-14).^22^ Our results suggest that full maturation of casp-3 may require the self-removal of the N-terminal peptide during extrinsic apoptosis. Additional studies employing a panel of cell lines, concentrations of **10**, and intrinsic and extrinsic apoptotic inducers will help elucidate the importance of casp-3 to the final stages of apoptosis.

**Figure 5.**
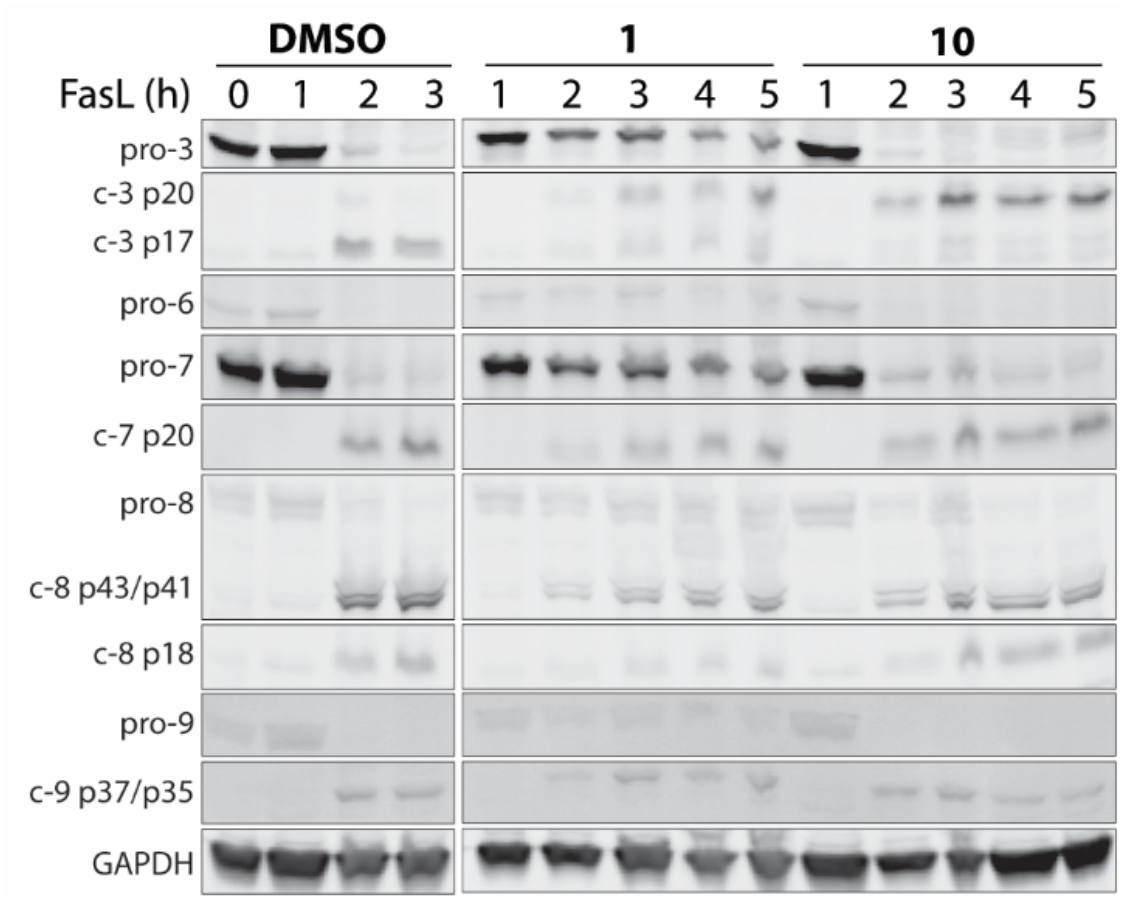
Profiling of caspase activation in extrinsically induced apoptotic Jurkat cells. Cells were treated with 20 μM of compound or (0.1% v/v) DMSO vehicle for 30 min before apoptosis was induced with 10 ng/mL MegaFasL. After the indicated time post-induction, cells were harvested and analyzed by Western blot. Caspases undergo rapid activation in DMSO (no inhibitor) control (Lanes 1-4). Treatment with **1** inhibits maturation of many caspases (Lanes 5-9). Treatment with **10** results in caspase profile mimicking the DMSO control, despite affording significant apoptotic protection (Lanes 1-4, 10-14).

## Conclusions

To date, the application of selective unnatural amino acid-based peptides designed for individual caspases were hampered by slow binding and thus limited their use to FRET-based substrates and activity probes in cellular studies. We present the design and validation of the first casp-3-selective peptide-based inhibitor, **10**, with the potency comparable to promiscuous DEVD-based inhibitors. Unlike previous generations of casp-3-selective inhibitors, **10** has the potency to affect biology and assist in cell-based studies. We demonstrate that aside from *in vitro* selectivity against other apoptotic caspases, **10** protects Jurkat cells from extrinsically induced apoptosis similarly to the general inhibitor Ac-DEVD-KE (**1**); however, treatment with **10** during apoptosis results in a caspase maturation profile comparable to untreated cells (e.g., DMSO vehicle control). This is in contrast to the promiscuous inhibitor 1, which generally suppresses the activation of all apoptotic caspases. Notwithstanding, an optimized chemical tool like **10**, will provide a critical resource to establish the intricacies and importance of individual caspases in programmed cell death, other biological processes, and disease. Additionally, we show that the negative charges of peptide-based molecules targeted to caspases do not require neutralization to be internalized by mammalian cells. This observation highlights the importance of carefully considering the effect of O-methylation with respect to solubility, permeability, potency, and esterase recognition when designing unnatural amino acid containing inhibitors for use in in biologically relevant cellular assays.

## Experimental Section

### Synthetic Protocols

Detailed protocols for the synthesis of the semicarbazide resin, AMAC-substrates and inhibitors are outlined in the Supporting Information available [insert hyperlink].^23–25^

### Preparation of recombinant caspases

Casp-3, casp-6, casp-7, casp-8, casp-9 and casp-10 were expressed and purified as previously described.^26^

### Enzymatic kinetic inhibition studies

Substrates, inhibitors and enzymes were diluted into assay buffer containing 20 mM PIPES, pH 7.4, 10 mM NaCl, 1 mM EDTA, 10 mM DTT and 10% sucrose. Solutions used to assay casp-9 and casp-10 were supplemented with 0.7 M sodium citrate. A 20 μL (2.5x) mixture of AMAC substrate (final concentration 100 μM; Ac-DEVD-AMAC for casp-3 and casp-7, Ac-VEID-AMAC for casp-6, and Ac-IETD-AMAC for casp-8, −9, and −10) and inhibitor was dispensed into a black 96-well Costar flat-bottom polystyrene plate. 60 μL (1.67 x) enzyme solution (final concentration: 1 nM casp-3, 25 nM casp-6, 5 nM casp-7, 25 nM casp-8, 100 nM casp-9, and 20 nM casp-10) was added, mixed for 10 seconds at 1000 rpm with fluorescence immediately read on a PerkinElmer EnVision plate reader with a λ_ex_ = [320] and λ_em_ = [460]. Data was analyzed using GraphPad Prism to determine k_inact_ and K_i_ in accordance with procedures previously outlined.^27^ However, progress curves were fit to 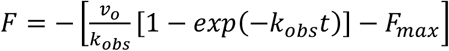 to account for the loss in fluorescence of AMAC substrates.

### X-ray crystallography of inhibitor-caspase complexes

Casp-3, and casp-7 were incubated at 250-300 μM with a two-fold molar excess of **8** or **10** in 20 mM Tris, pH 8.0, 150 mM NaCl, 10 mM DTT and 0.02% NaN_3_ for 2 h at 298 K and used immediately for co-crystallization experiments.

Crystals of casp-3 in complex with **8** were grown at 295 K after 1:1 dilution with 0.10 M sodium citrate, pH 5.4, 15.2 *%* PEG6000, 10 mM DTT, 0.02 *%* NaN_3_. Crystals of casp-3 in complex with **10** were grown at 295 K after 1:1 dilution with 0.10 M sodium citrate, pH 5.0, 13.6 % PEG6000, 10 mM DTT, 0.02 % NaN_3_. Crystals of casp-7 in complex with **8** were grown at 295 K after 1:1 dilution with 0.15 M sodium citrate, 1.6 M sodium formate, pH 5.0. Crystals of casp-7 in complex with **10** were grown at 295 K after 1:1 dilution with 0.15 M sodium citrate, pH 5.0, 1.6 M sodium formate. His_6_-tags were not removed as proteins crystallized readily.

Data was collected on single, flash-cooled crystals at 100 K with a cryoprotectant consisting of mother liquor and 20% PEG400 (casp-3) or 25% glycerol (casp-7) and were processed with HKL2000 in space groups I222 (casp-3) and P3221 (casp-7).^28^ The asymmetric unit for casp-3 contained one monomer of the biologically relevant homodimer and the complete homodimer for casp-7. X-ray data was collected to 1.50, 1.50, 2.65 and 2.35 Å resolution for casp-3:**8**, casp-3:**10**, casp-7:**8** and casp-7:**10**, respectively on beamline 9.2 at the Stanford Synchotron Radiation Lightsource (SSRL) (Menlo Park, CA). Data collection and processing statistics are summarized in Table S3.

### Structure solution and refinement

All caspase structures were determined by molecular replacement (MR) with Phaser ^29, 30^ using previously published casp-3 (4JJE) and casp-7 (4JJ8) as the initial respective search models. All structures were manually built with Coot^31^ and iteratively refined using Phenix ^32^ with cycles of conventional positional refinement. For the high-resolution casp-3 co-complex structures, anisotropic B-factor refinement was included. For the low-resolution casp-7 structures, non-crystallography symmetry restraints between the two subunits of the homodimer were applied during refinement. For all structures, the electron density maps clearly identified ligands were covalently attached to the caspase active-site cysteine. Water molecules were automatically positioned by Phenix using a 2.5 σ cutoff in *fo* – fc maps and manually inspected. For **8** co-complex structures with casp-3, the final R_cryst_ and R_free_ are 21.2% and 17.1% respectively; with casp-7, the final Rcryst and R_free_ are 19.7% and 24.4% respectively. For **10** co-complex structure with casp-3, the final R_cryst_ and R_free_ are 14.0% and 16.6% respectively; with casp-7, the final R_cryst_ and R_free_ are 19.6% and 23.5% respectively. All models were validated with the PDB server prior to deposition and all residues for the structures are located in the most favorable and additionally allowed regions in the Ramachandran plot. Coordinates and structure factors have been deposited in the Protein Data Bank with accession entries 6CKZ (casp-3:**8**), 6CL0 (casp-3:**10**), 6CL1 (casp-7:**8**), 6CL2 (casp-7:**10**). Structure refinement statistics are shown in Table S3.

### Cell Culture

Jurkat A3 cells were cultured as described by ATCC using 10% FBS and pen/strep/glutamine at 37 °C with 5% CO_2_. For all experiments, Jurkat cells were grown to near confluence, harvested and resuspended in fresh media to the indicated density. Apoptosis was induced using 10 ng/mL MegaFasL (AdipoGen^®^ Life Sciences).

### Apoptotic protection assays

Cells were plated into sterile Nunc Edge 96-well tissue culture-treated plates at a density of 10,000 cells/well and preincubated (or chased) with 20 μM of the indicated inhibitor or DMSO vehicle for the indicated time before (or after) apoptotic induction. 3 h after induction, cellular viability was measured using CellTiter-Glo^®^ on a PerkinElmer EnVision plate reader.

### PARP cleavage in apoptotic Jurkats

Jurkats were resuspended to 2 × 10^6^ cells/mL and treated with inhibitor or DMSO vehicle (30 min, 37 °C, 5 % CO2) before apoptosis was induced. After 3 h, the cells were harvested, washed twice with an equal volume of ice-cold PBS and resuspended at 25 × 10^6^ cells/mL with non-denaturing lysis buffer supplemented with protease inhibitors (Pierce A32965). Samples were then nutated (4 °C, 30 min), clarified (10 min, 21,000 × *g*), and analyzed by Western blot.

### Caspase activation studies

Jurkat cells were resuspended to 2 × 10^6^ cells/mL and treated with inhibitor or DMSO vehicle (30 min, 37 °C, 5 % CO2) before apoptosis was induced. At the indicated time point, cells were harvested, resuspended in CHAPS cell extract buffer (CST 9852), lysed by freeze-thaw (3 cycles), clarified (10 min, 21,000 × g), and analyzed by Western blot.

### Western blot analysis

Proteins were resolved by SDS-PAGE, transferred to PVDF membranes, blocked with 5% non-fat dry milk in TBST (0.1% Tween-20) and probed with the indicated antibodies. The primary antibodies were used as follows: anti-GADPH (Cell Signaling 2118, 1:2000), anti-PARP (Cell Signaling 9542, 1:1000), anti-pro/caspase-3 (Cell Signaling 9668, 1:1000), anti-pro/caspase-6 (Cell Signaling 9762 1:1000), anti-pro/caspase-7 (Cell Signaling 9492, 1:1000), anti-pro/caspase-8 (Cell Signaling 9746, 1:1000), anti-pro/caspase-9 (Cell Signaling 9502, 1:1000). Blots were incubated with primary antibodies (4 °C, overnight), washed (rt, 3 × 5 min, TBST) and incubated with secondary antibodies (LICOR IR800CDW or IR680RD, 1: 10,000) for 1 h at ambient temperature. Blots were further washed (3 × 5 minutes, TBST) and visualized on a LICOR Odyssey Scanner.

## Supporting Information

Detailed information about synthetic procedures and x-ray crystallography statistics (PDF).

## Author Contributions

A.S. and D.W.W. conceived of the project. A.S. and G.E.G.P. prepared the caspase proteins. A.S. performed the crystallography and structure analysis; *in vitro* kinetics studies and analysis; synthesized all peptides, substrates, and peptide-based inhibitors; and performed all cellular assays. The manuscript was written through contributions of all authors. All authors have given approval to the final version of the manuscript.

## Funding Sources

The authors gratefully acknowledge financial support from The Scripps Research Institute and the U.S. National Institutes of Health 5R21AI112796 and 1R01GM118382 (to D.W.W.).

## Notes

The authors declare no competing financial interests.

## ACKNOWLEDGMENT

We thank I. Wilson, R. Stanfield, M. Elsliger, and X. Dai for x-ray data collection and computational assistance, K. Backus for helpful discussions, H. Rosen and R.L. Wiseman for access to instrumentation, and the staff of the Stanford Synchrotron Radiation Lightsource.

