## Supplemental Information for "A selective and rapid cell-permeable inhibitor of human caspase-3"

### Table of Contents

**S2** Figure S1: Naïve $f_o$-$f_c$ densities of **8** and **10** in complex with casp-3 and casp-7

**S3** Figure S2: Interaction schematics of **8** and **10** in complex with casp-3 and casp-7

**S4** Table S1: X-ray data and structure refinement for all co-complex structures

**S5** Experimental section with general methods

**S8** Supporting Information references

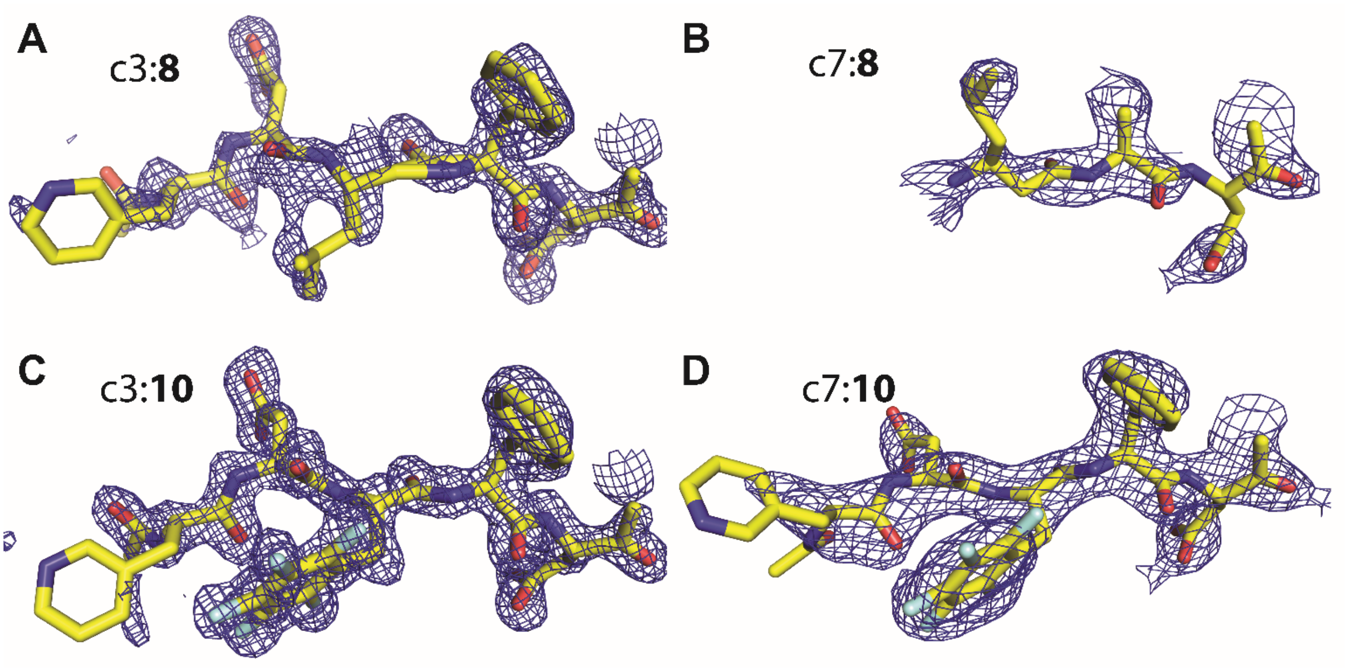


**Figure S1.** Related to Figure 2; Naïve $f_o$-$f_c$ densities of **8** and **10** in complex with casp-3 and casp-7. (A) casp-3:**8** (B) casp-7:**8** (C) casp-3:**10**(D) casp-7:**10**. Density for residues is weaker when bound to casp-7.

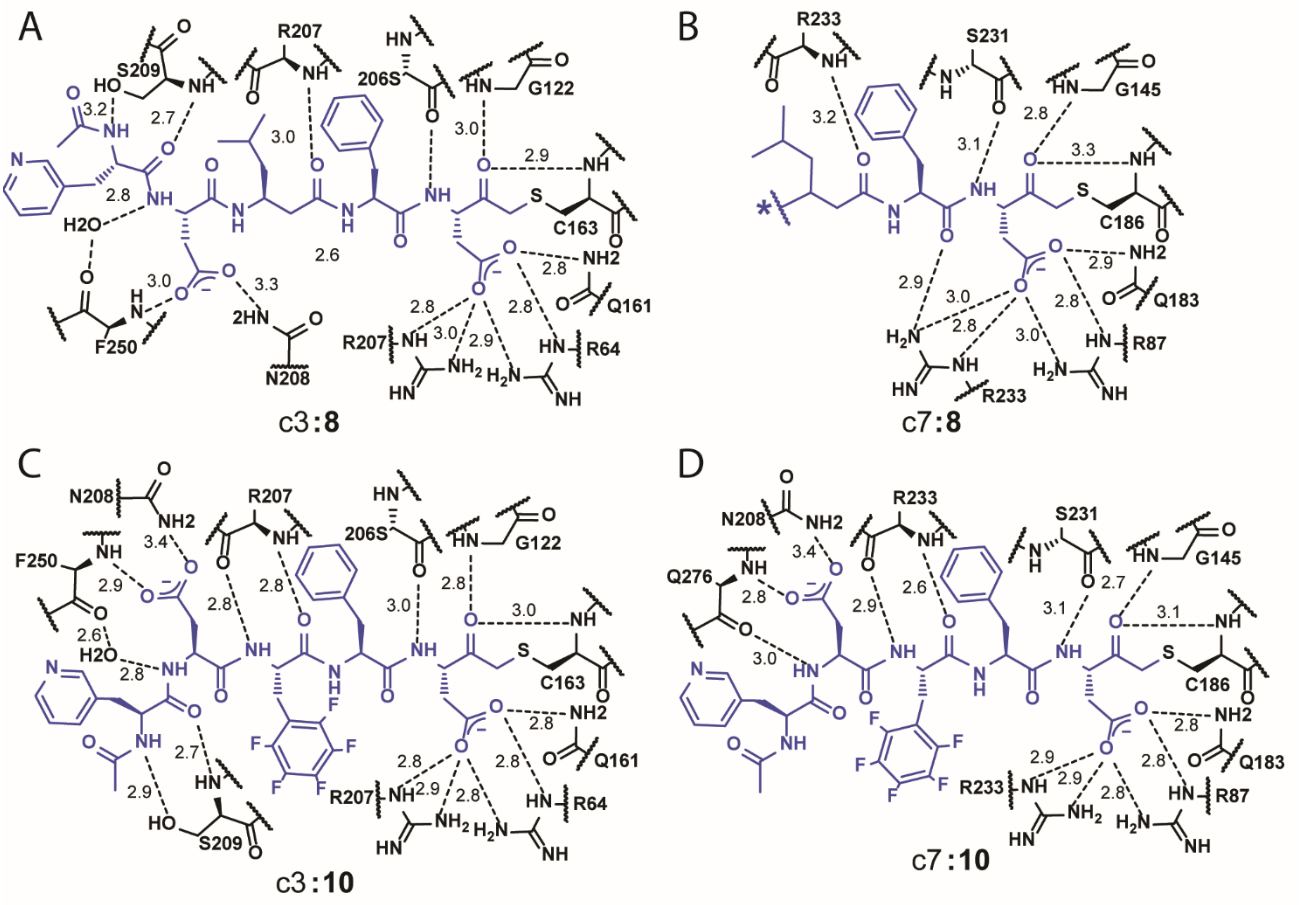


**Figure S2.** Related to Figure 3; Interaction schematics of **8** and **10** in complex with casp-3 and casp-7. (A) casp-3:**8** (PDB ID: 6CKZ) (B) casp-7:**8** (PDB ID: 6CL1) (C) casp-3:**10** (PDB ID: 6CL0) (D) casp-7:**10** (PDB ID: 6CL2).

**Table S1.** Related to Figure 3; caspase-3 co-complex X-ray data processing and structure refinement statistics

| Structure | casp-3:**8** | casp-3:**10** | casp-7:**8** | casp-7:**10** |
|---|---|---|---|---|
| PDB ID | 6CKZ | 6CL0 | 6CL1 | 6CL2 |
| Space group | I222 | I222 | $P3_221$ | $P3_221$ |
| Unit Cell Parameters (a,b,c) (Å) | 67.1, 83.9, 95.9 | 67.1, 84.1, 96.4 | 88.4, 88.4, 186.4 | 88.3, 88.3, 186.5 |
| Unit Cell Angles (z,y,z) | 90, 90, 90 | 90, 90, 90 | 90, 90, 120 | 90, 90, 120 |
| Data Processing | | | | |
| Resolution range (Å) (outer shell) | 50.0-1.50 (1.53-1.50) | 50.0-1.50 (1.53-1.50) | 50.0-2.65 (2.70-2.65) | 50.0-2.35 (2.39-2.35) |
| Unique reflections | 41,773 (1,635) | 40,755 (1,309) | 25,060 (1,232) | 35,850 (1,783) |
| Completeness (%) | 95.8 (75.8) | 92.6 (60.2) | 99.3 (99.8) | 99.8 (99.5) |
| Redundancy | 6.3 (4.6) | 7.6 (6.7) | 7.6 (7.5) | 8.1 (6.7) |
| $R_{meas}$ (%)[1] | 6.9 (50.9) | 8.8 (83.8) | 15.9 (112.6) | 17.0 (121.1) |
| $R_{merge}$ (%)[2] | 5.1 (35.1) | 6.7 (79.0) | 5.7 (40.7) | 12.9 (83.7) |
| $R_{p.i.m.}$ (%)[3] | 2.7 (22.8) | 3.1 (31.4) | 12.2 (91.0) | 5.9 (46.2) |
| Average I/Average σ(I) | 28.2 (2.6) | 28.6 (3.9) | 16.7 (2.5) | 14.9 (1.5) |
| $CC_{1/2}$ | 97.4 (88.9) | 97.0 (85.1) | 95.1 (81.9) | 93.3 (70.4) |
| Refinement | | | | |
| Resolution range (Å) | 41.9-1.50 (1.54-1.50) | 42.1-1.50 (1.53-1.50) | 48.2-2.65 (2.77-2.65) | 48.2-2.35 (2.41-2.35) |
| No. reflections[4] (test set) | 41,762 (2,101) | 40,752 (2,057) | 25,048 (1,189) | 35,790 (1,921) |
| $R_{cryst}$ (%)[5] | 15.3 (21.2) | 14.0 (17.4) | 19.7 (28.8) | 19.6 (27.3) |
| $R_{free}$ (%)[5] | 17.14 (25.5) | 16.4 (18.9) | 24.4 (36.8) | 23.5 (28.8) |
| Protein atoms / Waters | 1,955 / 206 | 2,059 / 250 | 3,625 / 28 | 3,642 / 74 |
| CV[6] coordinate error (Å) | 0.13 | 0.11 | 0.35 | 0.24 |
| Rmsd bonds (Å) / angles (°) | 0.007 / 1.99 | 0.017 / 1.523 | 0.009 / 1.224 | 0.007 / 0.954 |
| B-values protein/waters/ligands (Å²) | 21 / 33 / 39 | 18 / 30 / 25 | 47 / 44 / 66 | 46 / 42 / 66 |
| Ramachandran Statistics (%) | | | | |
| Preferred | 98.7 | 98.0 | 97.6 | 97.8 |
| Allowed | 1.3 | 2.0 | 2.4 | 2.2 |
| Outliers | 0.0 | 0.0 | 0.0 | 0.0 |

[1)] $R_{meas} = \{\Sigma_{hkl}[N/(N-1)]1/2\Sigma_i|I_{i(hkl)} - <I_{(hkl)}>|\} / \Sigma_{hkl}\Sigma_i I_{i(hkl)}$, where $I_{i(hkl)}$ are the observed intensities, $<I_{(hkl)}>$ are the average intensities and N is the multiplicity of reflection hkl. [2)] $R_{merge} = \Sigma_{hkl}\Sigma_i|I_{i(hkl)} - <I_{(hkl)}>| / \Sigma_{hkl}\Sigma_i I_{i(hkl)}$ where $I_{i(hkl)}$ is the $i^{th}$ measurement of reflection h and $< I_{(hkl)}>$ is the average measurement value. [3)] $R_{p.i.m.}$ (precision-indicating $R_{merge}$) = $\Sigma_{hkl}[1/( N_{hkl} - 1) ]^{1/2}\Sigma_i|I_{i(hkl)} - <I_{(hkl)}>|/\Sigma_{hkl}\Sigma_i I_{i(hkl)}$. [4)] Reflections with I > 0 were used for refinement.[1-3] [5)] $R_{cryst} = \Sigma_h||F_{obs}|-|F_{calc}||/\Sigma|F_{obs}|$, where $F_{obs}$ and $F_{calc}$ are the calculated and observed structure factor amplitudes, respectively. $R_{free}$ is $R_{cryst}$ with 5.0% test set structure factors. [6)] Cross-validated (CV) Luzzati coordinate errors.

### Experimental Section

- General methods
- General synthetic scheme for Fmoc-Asp(OtBu)-KE, semicarbazide resin (SCR) and coupling
- Synthesis of acyloxymethyl ketone inhibitors
- General synthetic scheme for 2-aminoacridin-9(10H)-one (AMAC), Fmoc-Asp(OtBu)-AMAC and loading onto 2-chlorotrityl resin

### General methods

All reagents and solvents were purchased from commercial suppliers and were used without further purification. All solution-phase reactions were performed in an inert atmosphere of dry nitrogen or argon. Reactions were monitored on TLC plates (silica gel 60, F254 coating, EMD Millipore, 1057150001), and spots were either monitored under UV light (254 mm) or stained with ninhydrin.

### General synthetic scheme for Fmoc-Asp(OtBu)-KE, semicarbazide resin (SCR) and coupling

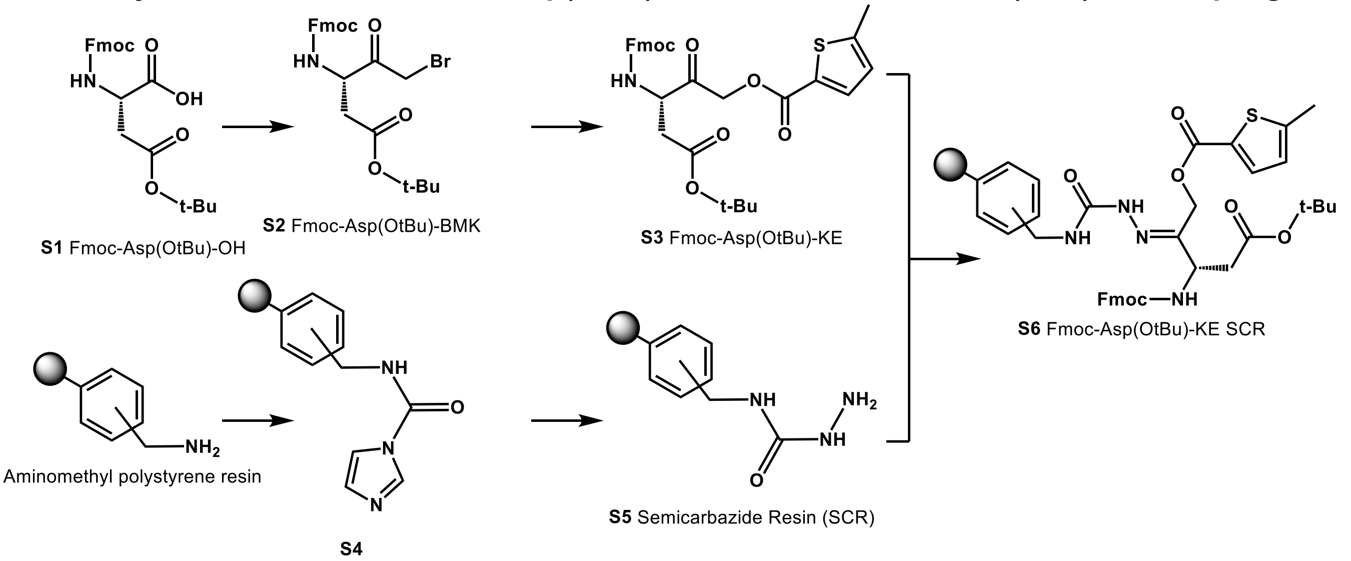


The following supplemental protocols are based on previously reported procedures used to generate cysteine protease inhibitors.[4-6]

### Synthesis of acyloxymethyl ketone inhibitors

*Fmoc-Asp(OtBu)-BMK,* **S2.** To a stirred solution of amino acid **S1** (4.12 g, 10.0 mmol) in DCM/ether at 0 °C within a 200 mL flame polished round bottom flask, N-methylmorpholine (1.06 g, 10.5 mmol) and isobutyl chloroformate (1.434 g, 10.5 mmol) were added dropwise. After 15 min, ethereal diazomethane was generated and distilled from Diazald® (6.43 g, 30.0 mmol) in accordance to procedures outlined in Aldrich Technical Bulletin AL-180 into stirred solution over the course of 30 minutes. After distillation, the reaction was allowed to warm to r.t. and continue for 1 hour. Glacial acetic acid was added dropwise after being chilled to quench excess diazomethane and 33% HBr in acetic acid added dropwise until a red tint persisted for more than 5 min. The solvent was removed *in vacuo*, redissolved in diethyl ether and subsequently washed with water, sat. aq. $NaHCO_3$ twice, sat. aq. NaCl, and dried over $MgSO_4$. Concentration *in vacuo* yielded **S2** in quantitative yield**,** which was used without further purification.
*Fmoc-Asp(OtBu)-KE,* **S3.** A flame dried 100 mL round bottom flask charged with 20 mL anhydrous DMF, anhydrous potassium fluoride (5.810 g, 100 mmol) and 5-methyl-2-thiophenecarboxylic acid (7.109 g. 50 mmol) was sonicated for 5 minutes. **S2**, dissolved in a minimal amount of anhydrous DMF, was added dropwise to stirred solution of carboxylic acid and base. After 30 minutes the solution was diluted with 250 mL EA, washed with 200 mL sat. aq. NaCl 2x, quickly with 1 M NaOH, sat. aq. $NaHCO_3$, sat. aq. NaCl, and dried over $MgSO_4$. Concentration *in vacuo* gave **S3** in 91% yield, which was used without further purification.

*Semicarbazide Resin (SCR),* **S5**. A flame-dried 500 mL round bottom flask charged with a magnetic stir bar, aminomethyl polystyrene resin (25 g, 28.75 mmol), N,N'-carbonyldiimidazole (46.62 g, 287.5 mmol) in 250 mL anhydrous DCM was stirred under positive argon pressure for 3 hr to yield **S4**. The resin was washed once with anhydrous DCM, once with anhydrous DMF, transferred into a new flame dried vessel and resuspended in 250 mL of anhydrous DMF. To this stirred solution, anhydrous hydrazine (55.29 g, 54.15 mL, 1725 mmol) was added gradually over 5 min. The reaction was stirred at room temperature for 1 hr to yield **S5**. The resin was filtered, washed with DCM 5x, MeOH 5x, dried thoroughly *in vacuo,* and stored at 4 °C.

*Fmoc-Asp(OtBu)-KE SCR,* **S6**. A flame-dried 100 mL round bottom charged with **S3** (5.00 g, 9.1 mmol) and **S5** (5.00 g, 1.15 mmol/g) was dried *in vacuo* for 6 hr and suspended in 50 mL anhydrous THF. This stirred solution was heated at 60 °C for 5 hr to yield **S6**. The excess amino acid derivative was recovered and the resin washed with DMF 2x, DCM 2x, MeOH 2x, dried thoroughly and stored at -20 °C.

*General solid phase peptide synthesis protocol to for inhibitors.* **S6** was presolvated in DMF for 30 min before two 15 min treatments of 5% diethylamine in DMF (1 mL/100 mg) to deprotect the N-Fmoc. Couplings were performed for 20 min using N-Fmoc-protected amino acids, HCTU, and DIPEA at a 3:3:10 ratio with respect to the loading of the resin, which was determined by Fmoc quantitation. Acetylation was performed using a 0.5 M solution of acetic anhydride and 10 eq. of DIPEA in DMF. The resin was washed with two aliquots of DCM and DMF in between each step. Prior to cleavage, the resin was additionally washed with DCM and MeOH 3x before being dried *in vacuo.* Cleavage was performed using 1 mL/100 mg resin of TFA/water/TIPS at a 95:2.5:2.5 ratio for 1 hr. The resin was washed with another aliquot of cleavage cocktail and the combined cleavage solutions were concentrated before precipitation with cold diethyl ether. The pellet was dried under a stream of argon and dissolved a minimal volume of DMSO before purification by preparatory reverse phase HPLC (19x100 mm XBridge C18, CH3CN/H2O/0.1% TFA, 10:90 to 90:10 over 15 minutes; 20 mL/min) and lyophilization.

**General synthetic scheme for 2-aminoacridin-9(10H)-one (AMAC), Fmoc-Asp(OtBu)-AMAC and loading onto 2-chlorotrityl resin**

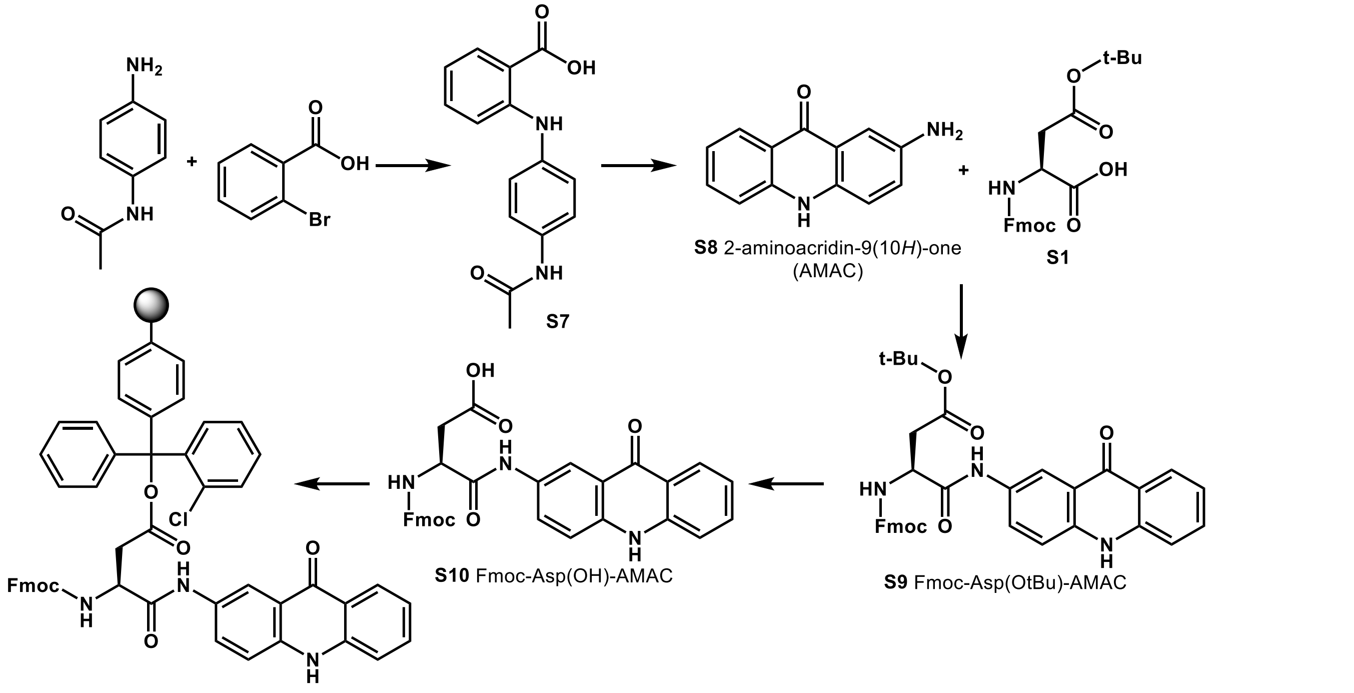


**S11** Fmoc-Asp(2-ClTrt)-AMAC

### Synthesis of AMAC substrates

*2-aminoacridin-9(10H)-one (AMAC),* **S8.** [7] A 100 mL round bottom flask was charged with 2-bromobenzoic acid (6.66 g, 33.1 mmol), N(4-aminophenyl)acetamide (5.00 g, 46.2 mmol), Potassium Carbonate (5.00 g 36.2 mmol), and copper (I) chloride (0.037.2 g, 0.38 mmol) in 25 mL isoamyl alcohol and refluxed at 155 °C for 4 hr. The solution was concentrated to near dryness *in vacuo* and diluted with EA, washed with 2 N HC l twice, washed with sat. aq. NaCl, dried over $MgSO_4$ and purified by silica on a 2 to 5 to 10 % MeOH gradient in DCM + 1% AcOH yielding 4.00 g of intermediate **S7.** A 100 mL round bottom flask charged with **S7 (**4.00 g, 14.8 mmol) in 30 mL concentrated sulfuric acid was refluxed at 100 °C for 3 hr. After cooling slightly, 60 g of ice was added and the solution was refluxed for another 2 hr to ensure complete deacetylation. The solution was chilled and neutralized to pH >7 with potassium hydroxide. The solid was filtered and washed with cold water and the product collected with MeOH. Removal of solvent *in vacuo* yielded **S8** in 26% yield over 2 steps and used without further purification.

*Fmoc-Asp(OH)-AMAC,* **S10.** [7]A 20 mL scintillation vial with 12 mL N-methyl-2-pyrrolidone (NMP) was sequentially charged with Fmoc-Asp(OtBu)-OH (0.741 g, 1.80 mmol), HBTU (0.682 g, 1.80 mmol), DIPEA (0.388 g, 3.00 mmol), and AMAC (0.315 g, 1.50 mmol) and then stirred for 2 hr at r.t. The product was precipitated by addition of 5% aq. $NaHCO_3$ (80 mL), filtered, and washed with cold water. The product was resuspended with MeOH, transferred, dried *in vacuo* and treated with 7.5 mL of a 95:2.5:2.5 TFA:water:TIPS cocktail for 1 hr at r.t. The cleavage solution was concentrated to less than 2 mL and the product precipitated by the addition of cold dry diethyl ether. The solid was washed with additional cold ether to give **S10** in 91% yield and used without further purification.

*Fmoc-Asp(2-ClTrt)-AMAC,* **S11**. A 60 mL fritted syringe charged with 2-chlorotrityl chloride resin (2.00 g, 1.1 mmol/g) was presolvated for 30 min with anhydrous DCM and washed with anhydrous DMF. **S10** (0.603 g, 1.37 mmol) and DIPEA (1.77 g, 13.7 mmol) in 10 mL anhydrous DMF was added and the vessel rocked for 90 min at r.t. to yield loaded resin, **S11**. The resin was washed twice with DCM and DMF before treatment with 20 mL of 20% pyrrolidine in DMF for 20 min to deprotect the N-Fmoc and cap unused resin sites. The resin was washed with DCM 2x and MeOH 2x before being dried thoroughly *in vacuo* and stored at -20 °C.

*General solid phase peptide synthesis protocol to synthesize substrates***.** Tetrapeptide fluorogenic substrates were synthesized similarly to peptidyl inhibitors except a single 20-min treatment with 20% pyrrolidine was used in place of two 15-min treatments with 5% DEA to deprotect N-termini. Table S1 lists the substrates used for each caspase analyzed and their corresponding $K_m$'s.
